# EukRef: phylogenetic curation of ribosomal RNA to enhance understanding of eukaryotic diversity and distribution

**DOI:** 10.1101/278085

**Authors:** Javier del Campo, Martin Kolisko, Vittorio Boscaro, Luciana F. Santoferrara, Ramon Massana, Laure Guillou, Alastair Simpson, Cedric Berney, Colomban de Vargas, Matt Brown, Patrick Keeling, Laura Wegener Parfrey

## Abstract

Environmental sequencing has greatly expanded our knowledge of micro-eukaryotic diversity and ecology by revealing previously unknown lineages and their distribution. However, the value of these data is critically dependent on the quality of the reference databases used to assign an identity to environmental sequences. Existing databases contain errors, and struggle to keep pace with rapidly changing eukaryotic taxonomy, the influx of novel diversity, and computational challenges related to assembling the high-quality alignments and trees needed for accurate characterization of lineage diversity. EukRef (eukref.org) is a community driven initiative that addresses these challenges by bringing together taxonomists with expertise spanning the complete eukaryotic tree of life and microbial ecologists that actively use environmental sequencing data for the purpose of developing reliable reference databases across the diversity of microbial eukaryotes. EukRef organizes and facilitates rigorous sequence data mining and annotation by providing protocols, guidelines and tools to do so.

## Introduction

Most lineages of eukaryotes (organisms with nucleated cells) are microbial, and eukaryotic diversity extends far beyond the familiar plants, fungi, and animals. Eukaryotic microbes—protists—include diverse lineages of mainly unicellular organisms, harboring a wide range of locomotion and trophic modes, including for example algae, heterotrophic flagellates, amoebae, ciliates, specialist parasites, and Fungi-like organisms among others. Although this term is polyphyletic, it was widely used for convenience to describe smallest size fraction eukaryotic communities in recent ecological studies, delineating them from bacteria and archaea. Collectively, they are important to ecological processes and to human health. Protists include important primary producers, particularly in aquatic ecosystems, as well as consumers that eat bacteria, algae, fungi, other protists, and even small metazoans, and thereby link microbial productivity to higher trophic levels [1]. Other lineages of protists recycle nutrients as decomposers or live as symbionts of other organisms. In fact, animals (including humans) are routinely colonized by eukaryotic microbes that run the gamut from parasites to commensals to mutualists [2].

Environmental sequencing efforts over the last 15 years [3,4] have greatly expanded the known extent of eukaryotic diversity, and the pace of data generation continues to grow. These efforts have identified many apparently novel lineages that have never been cultivated, and have transformed our understanding of the environmental distribution of numerous taxa [5]. The majority of environmental sequence data is based on the small subunit ribosomal DNA (also called 18S rDNA) because it is universally present, has been sequenced for the most comprehensive array of known taxa, and has a combination of conserved regions for primer design and variable regions that enable taxon identification [6]. With the advent of high-throughput sequencing, millions of sequences from hundreds of microbial communities covering a huge diversity can now be rapidly characterized within a study, enabling a broader community of researchers without a strong taxonomic background to investigate the temporal dynamics [7] and the spatial distribution of eukaryotic taxa within [8,9] and across [10,11] ecosystems, to test hypotheses about how eukaryotic communities are structured and how they respond to environmental change.

## Building a better database

Environmental sequencing may be transformative in all these ways, but these molecular datasets are only as good as the reference database used to annotate the data—at least for the multitude of studies that aim to assess eukaryotic diversity in a given ecosystem. Reference databases of ribosomal DNA bring together sequences from known isolates as well as Sanger-sequenced environmental datasets. The two main databases for eukaryotic ribosomal DNA sequences, SILVA [12] and the Protist Ribosomal Reference Database (PR2) [13], have improved dramatically in recent years, but substantial challenges remain. Views on eukaryotic relationships have changed in recent years [14]. Existing databases (PR2, SILVA, and the International Nucleotide Sequence Database Collaboration or INSDC [15]) differ in terms of numbers of sequences, taxonomic annotations, number of taxonomic ranks, and even inclusion of major lineages of eukaryotes [16] (Figure 1A). Thus, the database used for annotation strongly influences the taxa and the taxonomic resolution reported in a study. As an example, we used three different databases to annotate the same sequences from the BioMarKs [9] dataset, and even at the highest taxonomic level (the rank below “Eukarya”) got very different pictures of the taxonomic composition of the survey (Figure 1B).

**Figure 1.**
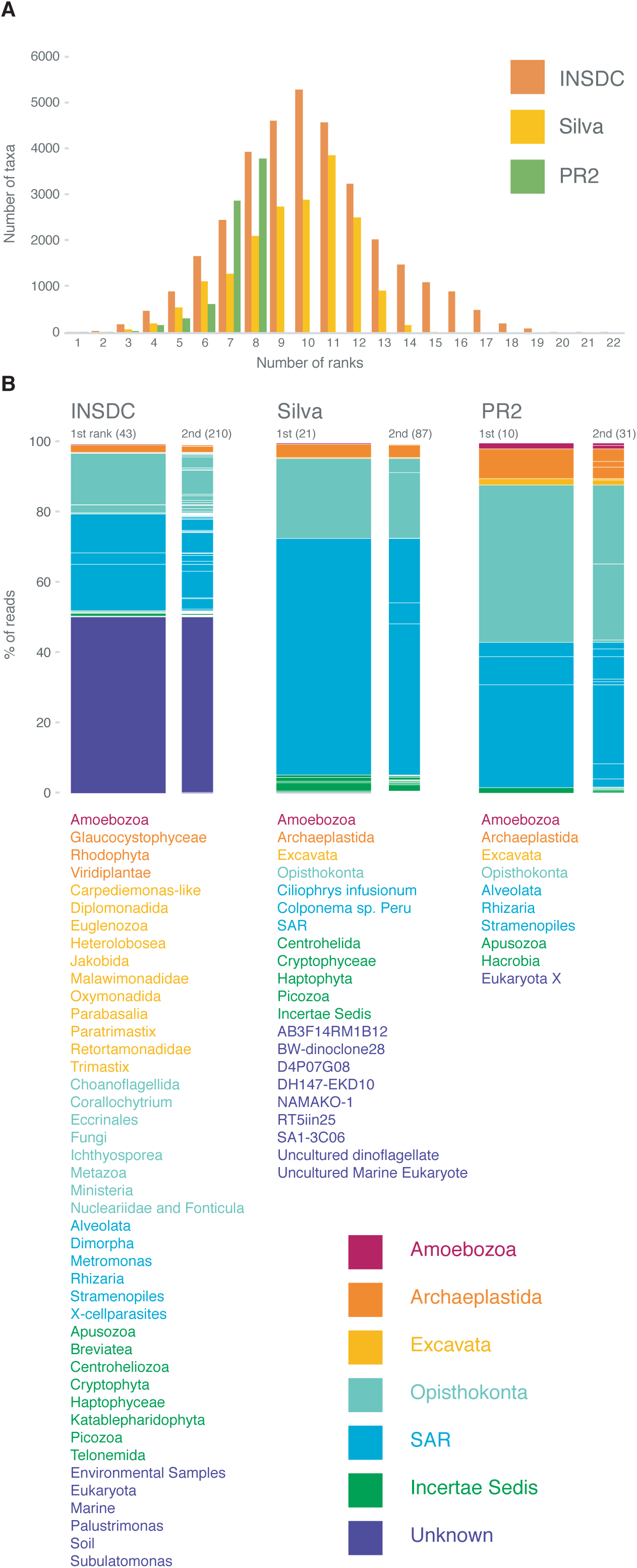
Databases comparison. (A) Distribution of the number of taxa used per rank in the most popular databases used as reference for protists metabarcoding analyses (INSDC GenBank release 215, Silva version 123.1, PR2 version 4.2) that we used to annotate the BioMarks dataset (B) Sequences from the BioMarKs environmental survey organized by the first (just below Eukarya) and the second taxonomic ranks of the three databases used. The bar plots show the main eukaryotic groups and how they are spitted into subgroups at each level. On top of each balrplot whiting brackets we show the number of taxa per rank. Below the first bar of each database the different taxa used in the taxonomic first rank. The taxa and the barplots are colored based on the eukaryotic super-groups defined by Adl et al. 2012 that represents the consensus taxonomy among the protistology community.

A subtler but equally important challenge is how to handle the variable taxonomic levels across clades of eukaryotes. For example, vertebrates have 15 taxonomic ranks in the INSDC taxonomy while the recently discovered Breviata lineage has only three, despite the fact that it diverged prior to the split between animals and Fungi [17]. This variability in ranks is sometimes a reflection of their known diversity, but nevertheless poses challenges during analysis. Many computational tools require a fixed number of ranks across taxa [23], and researchers generally want to be able to compare diversity across clades at a roughly equivalent level of diversity (or divergence time). Ideally, databases should flexibly handle several taxonomic ranks in a way that enables researchers to use standardized levels when necessary. Finally, the influx of vast amounts of data from environmental sequencing continues to reveal new lineages, and these data should ideally inform refinements in taxonomy and be incorporated into reference databases, so that new studies do not “rediscover” the same lineages, but rather refine what we already know about their diversity.

While ribosomal DNA is invaluable for taxonomic classification, it alone is unable to reliably disentangle deep eukaryotic relationships and is most powerful when combined with insight from multigene analyses (including phylogenomics) and/or morphological information. To address this, a flexible curation approach that can incorporate expert knowledge as well as insight from multigene molecular and morphological studies is supported by EukRef. This approach differs from that used by SILVA and Greengenes [18], which either rebuild the ribosomal tree for the whole dataset from scratch (Greengenes) or insert sequences into an existing and constant alignment and tree (SILVA). The use of backbone constraints based on published and robustly established relationships from phylogenomics and morphology facilitate incorporation of this knowledge and will be helpful in cases where this is warranted by existing data. For example, Fungi is a very well-established group of eukaryotes, but appears as polyphyletic in ribosomal DNA trees without backbone constraints [19]. EukRef guidelines recommend a tiered approach assessing multiple analyses to compare phylogenetic structure with and without constraints to understand their impact. In addition, we should be aware that novel lineages may have new insertions in ribosomal DNA or other differences that require rebuilding of the alignments. The improved reference database and phylogenetic tree will in turn enable better annotation of subsequent high-throughput sequencing studies.

## The EukRef curation process

The EukRef initiative proposes a platform where experts share the same guidelines and tools for the curation of taxonomic groups, with the fruits of these efforts to be reinvested into public databases. The initial phase of EukRef consists of development, coordination of experts, and yearly curation workshops, and is done in partnership with UniEuk, a network coordinating a taxonomic framework for eukaryotes. Here we presented the standardized guidelines and open source operational tools developed by this initiative after three years of existence, also available through EukRef.org. The final outputs of EukRef for each group are 1) a phylogenetic reference tree and alignment, 2) a curated reference database with accession numbers, curated classification string, and curated metadata, and 3) a list of sequences known to be problematic (such as chimeras).

To enable efficiency and consistency, the EukRef pipeline was developed to curate and annotate diverse eukaryotic lineages by researchers that are experts in that group to comprehensively capture existing sequence diversity for that lineage (Figure 2). Curation starts with a broadly sampled phylogenetic tree and alignment, and this initial set of sequences becomes the input to the EukRef workflow. The first step is an iterative retrieval of sequences from GenBank (INSDC) by BLAST [20] using a similarity threshold defined by the user depending on the targeted lineage. During this step sequences shorter than 500 bp and automatically detected chimeras using vsearch [21] are excluded. The expanded set of sequences retrieved from GenBank, together with the input sequences and relevant outgroups, are then aligned using MAFFT [22], automatically trimmed using trimAl [23], and a phylogenetic tree is constructed with RAxML [24]. The resulting tree is the starting point for curation and classification of sequences. Curators then manually examine the tree to identify discrepancies, such as long branches, which may be potential artifacts or chimeras that escaped the initial filtering, that should be removed (Figure 2). Following this cleaning step, a new alignment and tree are constructed with the remaining sequences. EukRef scripts then use the GenBank accession numbers for these sequences to retrieve the classification string and relevant metadata from GenBank and organize this information in a tab delimited file. This information, together with the tree, are the starting point for classification of the lineage and each sequence. These outputs are combined with previous taxonomic knowledge and improved metadata is manually incorporated throughout this process.

**Figure 2.**
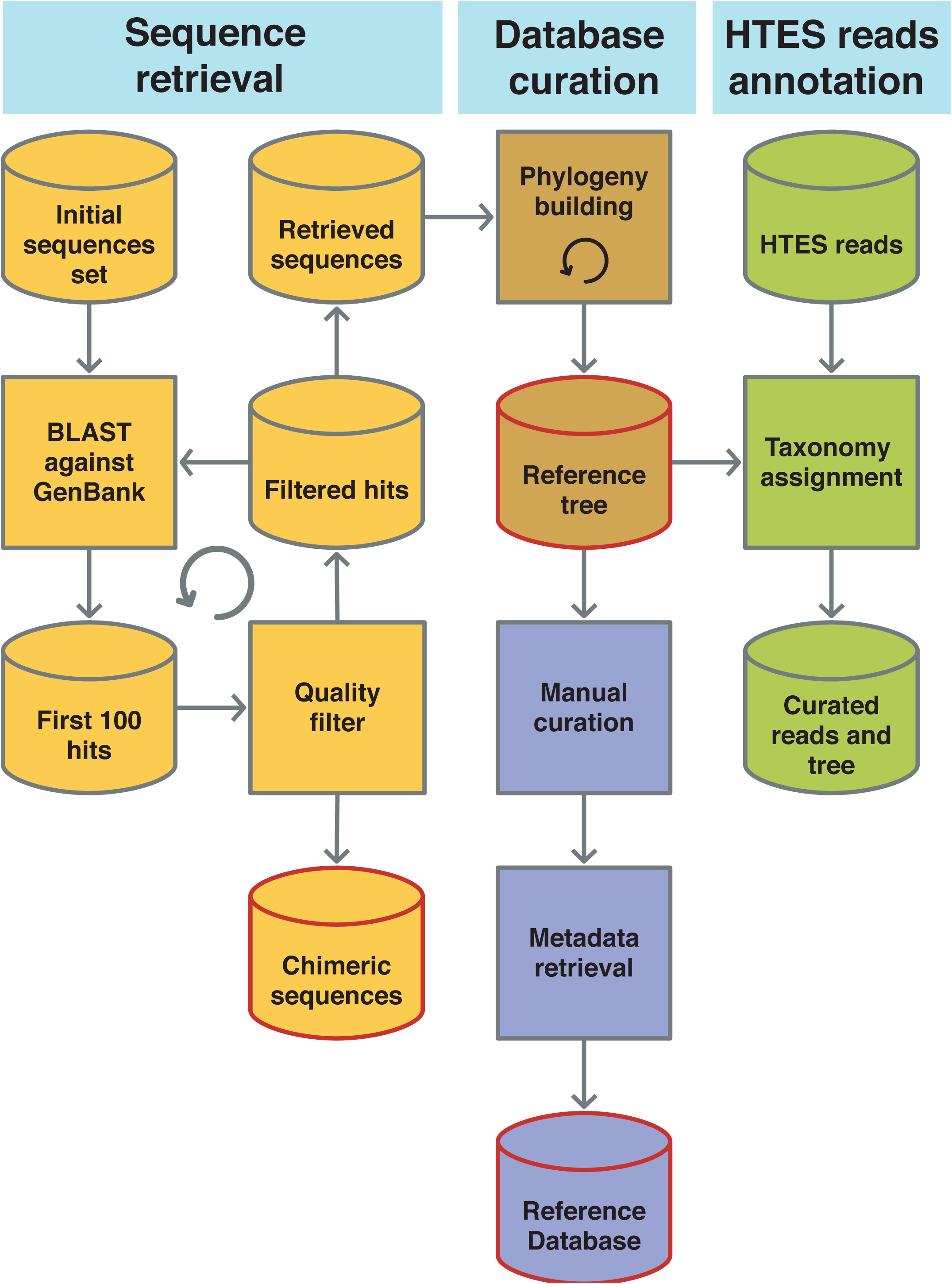
EukRef pipeline. Simplified scheme of the EukRef workflow with the outputs highlighted in red.

As a community, we have established guidelines for annotating sequences based on the phylogenetic tree and classification as part of the curation process, including guidelines for naming environmental clades. The curation process brings previous literature and expert knowledge to bear in annotating clades on the 18S tree generated in the EukRef pipeline, but presently informal names are largely ad hoc, especially those referring to environmental sequence clades; some degree of consistency would be both simpler and more informative. The proposed annotation guidelines are designed to be practical, stable, and compatible with downstream analyses that will use the curated databases. We recommend a conservative approach that minimizes the introduction of new names by relying on published literature and only assigning names to supported clades. Summarized guidelines are listed below (Box 2). Detailed guidelines and examples can be found at eukref.org/classification-guidelines. The guidelines for consistent naming of novel environmental clades should prove particularly useful: attaching a name is a key first step forward in scientific communication permitting the understanding of extent of diversity and mapping distribution of novel clades by allowing other scientists to recognize when they have found the same clade. Current ad hoc naming of these clades makes for substantial confusion when different arbitrary names are assigned to the same lineage, as inevitably happens.

Our approach also provides tools for attaching biological and environmental information to each sequence in the curated database, including basic habitat information, and whether a sequence came from a culture or morphologically identified isolate, or an environmental survey. Host associations are reported in the case of host-associated lineages (Box 3). To make this tool, we adopted standardized metadata annotation: MIxS (Minimum Information about any (x) Sequence) [25] and EnvO (Environment Ontology) [26]. The EukRef pipeline automatically assembles the complete set of sequences from NCBI associated with the clade of interest, but manual curation is required to vet the resulting phylogenetic tree, classify sequences, and transform the free-text retrieved from GenBank into MIxS and EnvO standard inputs. Additionally, we recommend the curators are encouraged to use relevant literature to fill in missing metadata and flesh out fields retrieved from GenBank to maximize the information attached to each reference sequence.

Altogether, these annotated datasets and the accompanying outputs are meant to provide a reliable tool for interpreting high-throughput sequencing surveys. The generated data will be available at the project website (eukref.org) and hosted long term in Dryad (datadryad.org). Each lineage-specific dataset will be integrated into the UniEuk [27] (unieuk.org) taxonomic framework implemented at EBI, and provide phylogenetic evidence for internal nodes and environmental clades of significance. In the long term these datasets will also be transferred to existing reference databases for eukaryotes, including SILVA, PR2, and eventually INSDC. The annotations will also be freely available to other databases that are currently taxonomically restricted, but might wish to expand to eukaryotes, such as the Ribosomal Database Project [28]. Ongoing curation and incorporation of newly available sequences will be facilitated by using Pumper [29], which allows an automatic sequence retrieval and tree building, and Sativa [30], which automatically annotates sequences in a tree. Both depend on the quality of the initial input, highlighting the need for high-quality initial annotation as implemented in EukRef.

## Ciliates as a case study

To illustrate both the curation process and why it is important, we annotated a well-known group of ciliates, the Heterotrichea, as a case study. The initial dataset imported into the pipeline consisted of only 9 SSU rDNA sequences published by Rosati et al. in 2004 [31] (Figure 3A). After 6 cycles of the sequence retrieval script, we obtained 412 sequences that were combined with outgroup sequences and used to build an initial tree (Figure 2). After discarding all the sequences that fall outside of the Heterotrichea, low quality sequences, and sequences less than 500 bp, we were left with 258 heterotrich sequences (Figure 3B) representing 37 OTUs clustered at 97% (Figure 3C). Of these 258 sequences 74% corresponded to isolates (previously cultured and/or isolated taxa), and 26% corresponded to sequences known only through environmental surveys. The classification of each of the 258 sequences was annotated based on the phylogeny, and the metadata curated using GenBank records and associated literature according to EukRef guidelines (Figure 3C). The classification was improved or corrected compared to the initial GenBank record for a quarter of the sequences, and metadata was added for the majority (70% of the sequences). In accordance with the EukRef guidelines, we have released the reference database, a list of short reads, a list of chimeras, an alignment, and a reference tree are publicly available at the EukRef website and Dryad (Supplementary Material). This example shows the efficiency of the EukRef pipeline to increase the taxon sampling and diversity coverage for this clade and collated available sequenced isolates as well as environmental sequences of previously unknown taxonomic affiliation. This increased taxon sampling and coverage, which in turn dramatically improved the phylogenetic resolution of the Heterotrichea. By curating basic environmental metadata, we also provided valuable information not only for the classification of this lineage, but also the ecology and environmental distribution of the Heterotrichea, as well as providing a tool to future researchers to more accurately annotate Heterotrichea reads from high-throughput environmental surveys.

**Figure 3.**
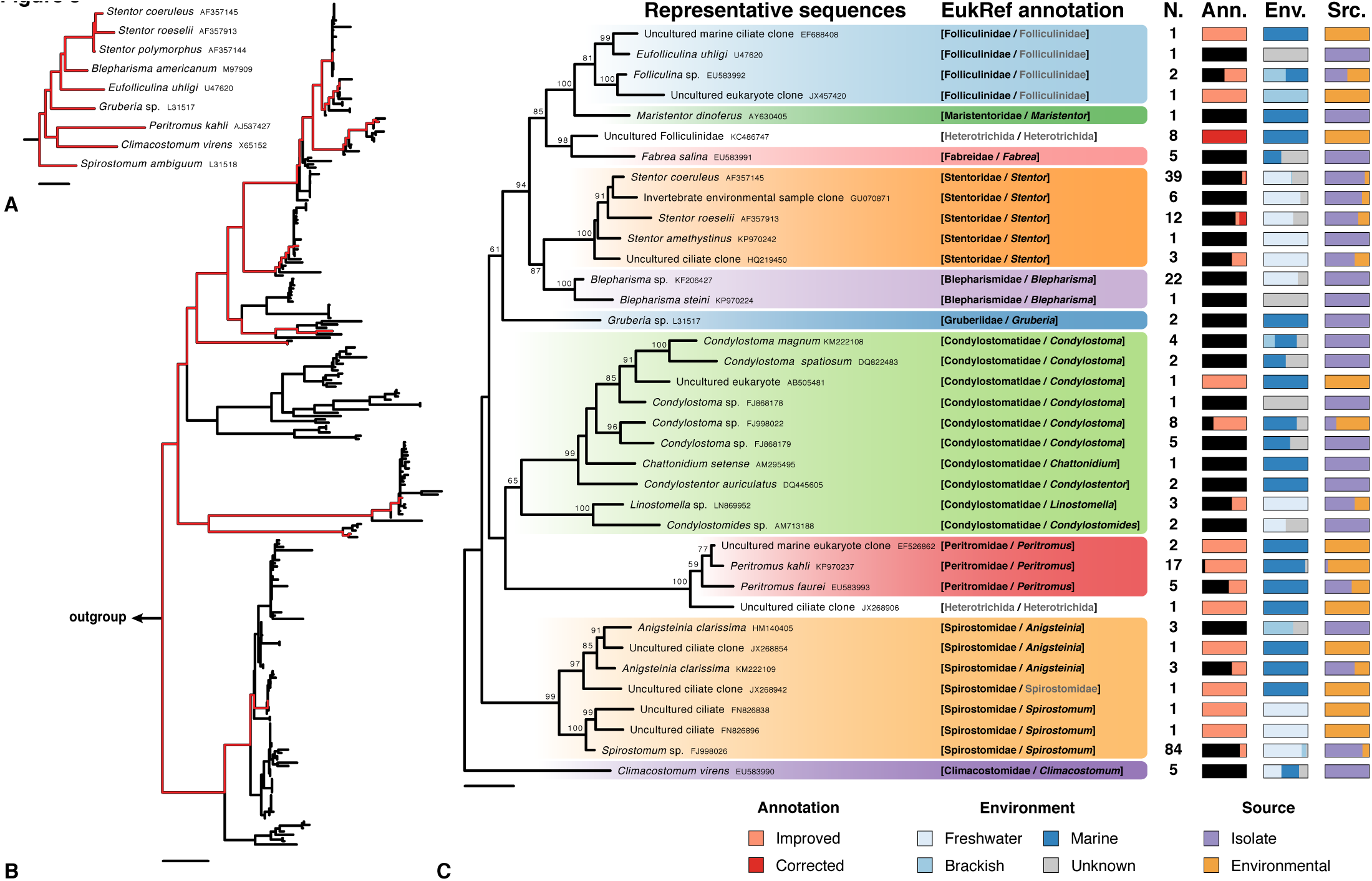
Case study: Heterotrichea, Ciliophora. A small set of sequences (A) is used as input to obtain related GenBank-deposited sequences that are then displayed in a phylogenetic tree (B). Representative sequences clustered at a 97% similarity threshold are used to obtain the final phylogenetic tree, which is used as a guide to perform the taxonomic annotation (C). Gaps in the unlimited-ranks classification are filled extending the annotation of broader taxa (examples are shown in grey). The classification is then propagated to all the sequences in the database (N., number per cluster). The percentage of sequences per cluster with an improved or corrected taxonomic annotation is shown in the *Ann.* column. Various metadata are manually appended to each sequence. As examples, environment and source of sequences are shown in columns *Env.* and *Src.*, respectively.

## Outreach and Training

Yearly weeklong intensive curation workshops organized in different parts of the world represent the core of the curation effort. These workshops bring together advisers (taxonomic experts) with curators—typically students and postdocs—that are actively investigating the taxonomy and diversity of a particular eukaryotic lineage to curate a reference database for a particular eukaryotic group that will further their research efforts.. The participants acquire the expertise for how to use the provided workflow and tools to gather and curate a ribosomal DNA database, and they gain experience working in a UNIX-based command line environment. The process of curating the classification and metadata for their retrieved sequences requires them to delve deeply into the literature for their lineage, improving their taxonomic knowledge of the eukaryotes, too. These early career scientists get also connected to the community of researchers studying protist classification and environmental distribution allowing them to expand their network and establish collaborations beyond the context of EukRef..

## Conclusions and future perspectives

EukRef brings together members of the community with expertise in the taxonomy of different eukaryotic lineages to curate available ribosomal DNA sequences from cultured isolates and morphologically identified organisms, as well as environmental surveys, all within in a phylogenetic framework. In the long term, EukRef and the associated UniEuk initiative aim to assemble a curated reference database of 18S rRNA gene sequences covering all eukaryotes. Taxonomists have the greatest knowledge of eukaryotic groups but are rarely involved in curating databases and seldom use existing environmental data. However, these are exactly the people needed to make sense of the vast diversity revealed in these studies. Bringing together taxonomists and microbial ecologists will provide better reference databases, which, our aim is toimprove the automatic annotation of eukaryotic environmental sequencing surveys being increasingly conducted by the broader research community.

## Source code

All source code for the EukRef pipeline is available from: https://github.com/eukref/eukref

## Acknowledgements

We thank to all of the EukRef workshop participants and advisors. We also thank Frederic Mahé and Alexey Kozlov for reviewing the scripts in their earliest versions. Thanks to the Gordon and Betty Moore Foundation, the National Science Foundation Division of Ocean Sciences, Biological Oceanography Cluster (Award 1545931, www.nsf.gov), and the International Society of Protistologists for their financial support. JdC, VB, and MK were supported by grants to the Centre for Microbial Diversity and Evolution from the Tula Foundation. JdC was supported by a Marie Curie International Outgoing Fellowship grant (FP7-PEOPLE-2012-IOF - 331450 CAARL). MK was supported by Fellowship Purkyne (Czech Academy of Sciences, Czech Republic) and by ERD Funds (CZ.02.1.01/0.0/0.0/16_019/0000759 CePaViP). LFS was supported by grant OCE1435515 from the U.S. National Science Foundation to G. McManus (University of Connecticut). CB was supported by the “Investissements d’Avenir” programme OCEANOMICS (ANR-11-BTBR-0008 OCEANOMICS). LWP is supported by an NSERC-DG.

## Boxes

## Box 1

Definitions as used by EukRef:

- Low level: less inclusive taxon (e.g., genus)
- High level: more inclusive (e.g., phylum)
- Database: Refers to tab delimited file constructed by EukRef curators containing information about the identity of a sequence, its classification, and environmental metadata.
- Clade: used here to refer to clade in a phylogenetic tree.
- Taxon: a group of organisms that has been assigned a name in previous literature (e.g., a genus, or a species).
- Group: a lineages or clade in a phylogenetic tree being curated.
- Chimera: DNA sequence that stem from two or more original sequences generated as a product of the DNA amplification process.

## Box 2

Classification Guidelines box:

1. Clades should be supported by previous literature and/or receive statistical support in the 18S phylogenetic tree to be named.
2. Use names that are established in the literature. These can by formal taxon names, informal names, or environmental sequence clade names.
3. EukRef uses named rankless levels (i.e., not adhering to Linnaean classification ranks) following Adl et al 2012. Use as many levels as needed.
4. Only annotate to the level for which there is support. Fill in blank ranks by propagating up from the lower levels (less inclusive) to higher levels.
5. Do not name clades that are not supported, or clades where the applicability of a name is ambiguous. (See website for examples and detailed guidelines)
6. Novel environmental clades may be named following the Novel Naming Environmental Clades guidelines (below).

Naming Environmental clades

- Only name lineages that are:
  - well-supported by bootstrap / posterior probabilities or possibly by clear 18S sequence signatures, and
  - composed of three or more clearly distinct sequence types, ideally from two or more different studies.
- Use a 3-5 capital letter code for the clade containing the environmental lineage. In most cases this should be the most inclusive clade being annotated (e.g., API for environmental clades within Apicomplexa). Avoid using different codes for each subclade. This introduces unnecessary names and instability because the position of environmental lineages often shifts in subsequent analysis.
- Number the lineages in some arbitrary order, for instance chronological order of their first appearance in a paper. (e.g., API3). Use numbers again after an underscore for sublineages (e.g., API3_2)
- Never re-use the same number—even if a lineage later disappears—to avoid confusion (e.g., MAST-5 no longer exists)
- Do NOT name isolated sequences, especially long-branches. These are potentially chimeras or bad-quality sequences. When isolated sequences look genuine (are not chimeras upon detailed inspection), they can be kept in the reference alignment and database because they may carry useful environmental information. These sequences should be identified simply by their clone name.

## Box 3

For the metadata annotation, we adopted standards from MIxS (Minimum Information about any (x) Sequence) and EnvO (Environment Ontology):

- source*: Does this sequence come from a culture or an environmental sequence.
- env_material: The environmental material level refers to the material that was displaced by the sample, or material in which a sample was embedded, prior to the sampling event. Environmental material terms are generally mass nouns.
- env_biome: Biome should be treated as the descriptor of the broad ecological context of a sample and defined by a certain biotic community and other factors environmental factors like climate.
- biotic_relationship: Life style, from free living to mutualistic symbiont.
- specific_host: For symbiotic lineages (including parasites). Host taxonomy ID (taxid) from NCBI.
- geo_loc_name: The geographical origin of the sample as defined by the country or sea name followed by specific region name.

**Supplementary Information 1**. Case study output 1, Heterotrichea alignment.

**Supplementary Information 2**. Case study output 2, Heterotrichea tree.

**Supplementary Information 3**. Case study output 3, Heterotrichea curated reference database.

**Supplementary Information 4**. Case study output 4, list of identified chimeras.

